# A novel algorithm to optimize generalized gamma distributed multiplicative noise with implications on speckle removal from OCT images

**DOI:** 10.1101/2020.10.07.329227

**Authors:** Divya Varadarajan, Caroline Magnain, Morgan Fogarty, David A. Boas, Bruce Fischl, Hui Wang

## Abstract

Optical coherence tomography (OCT) images are corrupted by multiplicative generalized gamma distributed speckle noise that significantly degrades the contrast to noise ratio (CNR) of microstructural compartments in biological applications. This work proposes a novel algorithm to optimize the negative log likelihood of the spatial distribution of speckle. Specifically, the proposed method formulates a penalized negative log likelihood (P-NLL) cost function and proposes a majorize-minimize-based optimization method that removes speckle from OCT images. The optimization reduces to solving an iterative gradient descent problem. We demonstrate the usefulness of the proposed method by removing speckle in OCT images of uniform phantoms with varying scattering coefficients and human brain tissue.

## I. Introduction

Optical coherence tomography (OCT) is an imaging technique that uses low temporal coherence light to obtain cross sectional images of an object at 1 - 20 *μm* resolution. The high resolution of OCT images and its ability to image in 3 dimensions make OCT an attractive imaging modality to study microstructure in biological tissue. Several recent studies have shown that optical properties of OCT images provide distinctive contrasts for anatomical landmarks [1]–[3] and clinical biomarkers that are valuable for pathological diagnosis and monitoring of disease progression and treatment outcome [4], [5].

The contrast in OCT images of biological tissue mainly reflects scattering events, which arise due to differences in refractive index, and backscattering events that result from various microstructural properties of the tissue compartments [6]. Each microstructure compartment attenuates the light propagating in the tissue with a rate defined by the scattering coefficient. Estimation of optical properties such as scattering coefficient in OCT images has been shown useful in numerous applications, including brain imaging, histology correlation, and pathology detection [1], [7], [8].

A common problem in contrast visibility and optical property estimation is the contamination of OCT images by speckle noise. Speckle is a form of multiplicative noise that occurs due to the interference of scattering waves from multiple regions of the microstructure [9], [10]. Destructive interference reduces the OCT signal, while constructive interference increases the intensity, giving speckle a high contrast appearance. Speckle significantly degrades the contrast-to-noise-ratio (CNR) between tissue structures and masks features that are similar in size to it, dramatically reducing the accuracy of quantitative analysis.

Speckle noise is typically eliminated by spatial averaging over multiple OCT acquisitions with uncorrelated speckle patterns [11], [12]. Since *ex vivo* fixed tissue lacks dynamic processes, the uncorrelated speckle patterns are created with different incident wavefronts, angular compounding, frequency compounding or combining polarization modes. In a recent study, we used the average of 50 percent overlapping tiles to reduce speckle noise and achieve large volumetric reconstruction of *ex vivo* human brain tissues [2], [3]. The resulting speckle contrast is inversely proportional to the square root of number of averages. However, this type of speckle reduction methods suffers from substantially increased acquisition time. For example, acquiring 50 percent overlapped data takes four times longer than acquiring non-overlap data.

A second class of techniques to remove speckle noise can be broadly referred to as post processing tools that apply denoising algorithms to the acquired OCT data. The high-contrast appearance of speckle has led to the usage of filtering-based methods [13]–[16] to suppress it. While filtering works well for homogeneous tissue, it distorts tissue boundaries and blurs structures, reducing the effective resolution of the image. Several additive noise-based denoising methods that denoise the logarithm of the OCT signal have been proposed. Most of these methods assume zero mean noise which does not hold for the log transformed speckle distribution, leading to a mean bias in the denoised signal [17]–[19]. Nevertheless, additive denoising methods based on non-local means (NLM) [20]–[22], wavelet transformation [23]–[25] and constrained least square methods [26]–[28] have been used to remove speckle. In addition, NLM approaches require repeated similar patches in an image to perform well [29] which can be problematic because human brain tissue is highly spatially varying between 1 - 20 *μm* resolution, wavelet methods can suffer from wavelet domain artifacts [29] and least squares fidelity is sub-optimal because it inherently and incorrectly assumes the speckle distribution to be log normal [9].

This brings us to the final class of maximum likelihood estimation based that directly optimize the spatial distribution of speckle. Speckle has been shown to be generalized gamma distributed by many previous studies [8], [30]–[35]. Several methods that either approximate the speckle distribution to Gaussian distribution [29], which does not model speckle accurately [9], or that optimize special cases of generalized gamma distribution such as Rayleigh [36]–[38], gamma [19], [39] and negative exponential [6], [40] distribution that might not generalize to all biological specimens [30], [35] have been proposed.

In this paper we propose a novel method that directly optimizes the generalized gamma distribution. The proposed method minimizes the negative log likelihood of the generalized gamma distribution penalized with a spatial regularization constraint using the majorize-minimize optimization method [41], [42]. We show that the overall minimization reduces to solving an iterative gradient descent procedure of convex cost functions. While we mainly use a quadratic spatial smoothness regularization function, the proposed framework is also flexible to accommodate other convex regularization constraints. We also show that our method theoretically generalizes to gamma and negative exponential distributions.

We compare the performance of our proposed method with the commonly used median filtering method and non local based Block Matching 3D (BM3D) denoising method [22] with simulations, phantom and human brain tissue experiments. We demonstrate the generalizability of our approach to remove speckle across multiple tissue types and multiple imaging resolution scales.

The paper is organized to briefly describe the background on speckle likelihood, derive the proposed algorithm called MM-despeckle, demonstrate speckle removal results, discuss implications of our approach and present conclusions.

## II. Background

### A. Maximum likelihood estimation (MLE) for denoising applications

MLE is a commonly used procedure to denoise data. The main assumption for MLE methods is that we can model the likelihood distribution *P_Y|X_*(*y; x*) of the acquired data. Broadly speaking, MLE methods find an estimate 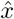 of the true signal *x* from the measured data *y* by maximizing the likelihood distribution. This is generally achieved by minimizing a penalized negative log likelihood (P-NLL) cost function,

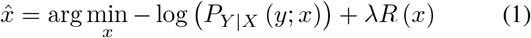

where *R*(*x*) is a regularization or penalty function and λ is the regularization parameter. The penalty is put in place to constrain the generally ill-posed nature of the negative log likelihood minimization. The next step involves finding an optimization procedure that minimizes the P-NLL cost function.

### B. Majorize-minimize Optimization Framework

In this work we specifically derive a procedure based on the majorize-minimize (MM) optimization framework [41], [42]. The MM procedure has been used in the past for denoising additive noise in magnetic resonance imaging [43]. The MM framework minimizes a complicated cost function indirectly by sequentially minimizing simpler convex functions that are tangential to and greater than or equal to the cost function. The tangential functions are referred to as majorants.

Mathematically, a majorant *G*(*x|x^i^*) of a cost function *C*(*x*), tangential to it at *x^i^* satisfies the following two criteria:

- The cost and the majorant meet only at a single point *x^i^*: *C*(*x^i^*) = *G*(*x^i^|x^i^*)
- The majorant function is greater than the cost function otherwise: *C*(*x*) < *G*(*x|x^i^*), ∀*x* ≠ *x^i^*

These conditions theoretically guarantee that the sequential minimization is monotonically decreasing the cost *C*(*x*) thereby minimizing it and that the majorants are convex functions thereby simplifying the minimization of each iteration.

### C. Speckle distribution in OCT images

Speckle in OCT images [6] is a form of multiplicative noise that is modeled at an arbitrary voxel *m* as

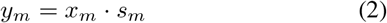

where *y_m_* is the measured OCT intensity, *x_m_* is the true OCT intensity and *s_m_* is the speckle noise. The generalized gamma based spatial distribution (GGD) of speckle 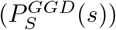 is represented mathematically as,

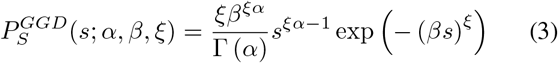

where Γ(·) is the gamma function, *α* and *ξ* are the shape parameters and *β* is the rate parameter. As an example, figure 1 shows an empirical distribution of speckle in an OCT image of a uniform scattering phantom with a scattering coefficient of 0.01/*μm*. The phantom was imaged by a 1300 *nm* spectral domain OCT at 3.5 *μm* isotropic resolution. The specifications of the OCT system and the scattering phantom are described in detail later in section IV. The histogram shows the intensities of a B-scan, which was normalized with the mean intensity of the corresponding depth. GGD fit of the histogram generated *α* = 1.14, *β* = 1.20 and *ξ* = 0.92. We used the source code from [44] for GGD fitting.

**Fig. 1.**
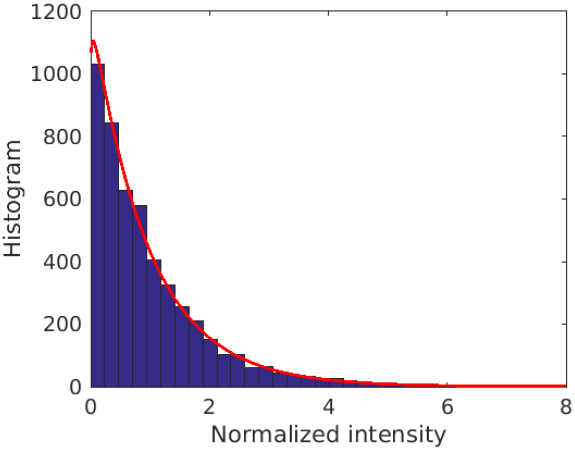
GGD fit for a uniform phantom: The figure plots the histogram of normalized intensity values representing speckle of a uniform phantom OCT image. The red curve is the generalized gamma fit to the histogram. GGD fits the OCT speckle well.

## III. III. Theory

### A. Negative log likelihood cost function

The likelihood of the OCT signal at a single voxel is described as the conditional probability of the measured intensity *y_m_* given the true signal *x_m_*. It can be derived from equations 1 and 2 as,

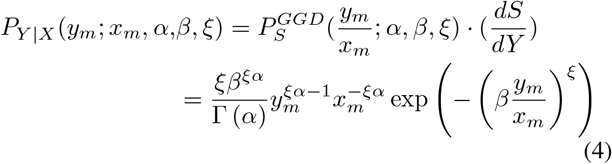

where we have substituted the derivative 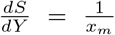, that followed from the multiplicative relationship of Eq. 1.

Assuming all voxels to be identical and independently distributed (i.i.d) samples of the likelihood distribution, the joint likelihood can be derived as,

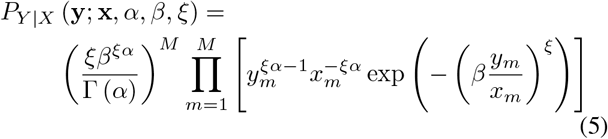

where M is the number of measurements (or voxels), m is the voxel index, y = [*y*_1_, ···, *y_M_*] ∈ *R^M^* is the vectorized measured OCT image of length M and x = [*x*_1_, ··· , *x_M_*] ∈ *R^M^* is the vectorized true OCT image of length M.

The P-NLL cost for speckle noise follows as,

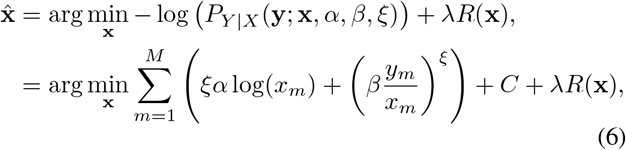

where 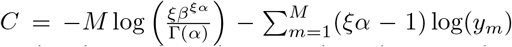 are constants that do not depend on x and can be ignored in the minimization.

### B. Majorant to P-NLL cost function

GGD NLL for an arbitrary voxel is a sum of a logarithm function *ξα* log(*x_m_*), and the power term 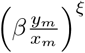 (see Eq. 7 and 8). For a positive valued true signal (*x_m_* > 0), the logarithm term is strictly concave with a negative second derivative, while the power term is strictly convex with a positive second derivative.

The tangential majorant M(x; x^*i*^) of GGD NLL is the sum of the convex power term and the tangent to the logarithm function, given by,

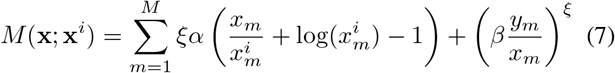

where the tangent at 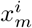 to log(*x_m_*) is given by 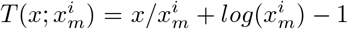.

Figure 2. plots the GGD NLL cost function in blue and shows the majorant function (*M*(*x; x^i^*)) at three points (*x^i^*) of the NLL cost function. As the minimum of the majorants moves from the red to the yellow and purple curve, the estimation gradually approaches the minimum of the NLL cost function.

**Fig. 2.**
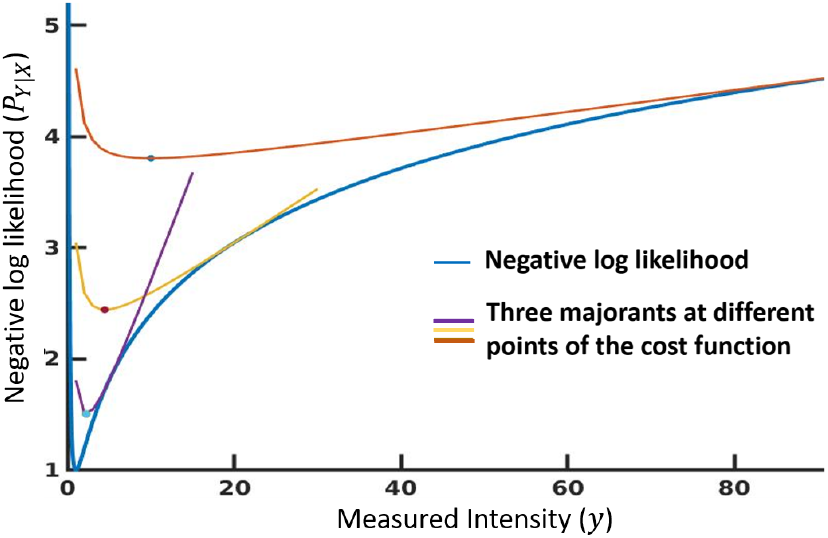
Example majorants of GGD. The figure plots NLL of a GGD with true intensity = 1 and majorants at three points on the NLL curve.

Assuming a convex regularization function, the majorant to the full P-NLL cost becomes

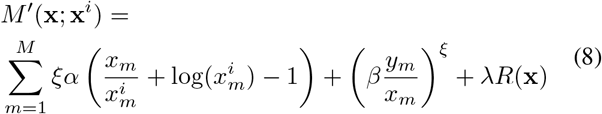

The likelihood distribution, P-NLL cost and majorant corresponding to special cases of gamma and negative exponential distributions have been derived in Supplementary Sec. I.

In this paper we use a spatial quadratic smoothness (Tikhonov) regularization function,

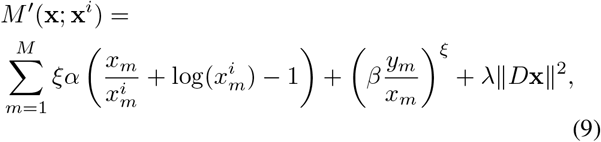

where *D* is the two dimensional finite difference matrix. The partial derivative at *x_m_* for the P-NLL majorant can be analytically solved as

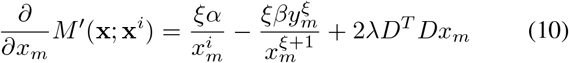

The majorant is minimized iteratively using gradient descent.

### C. MM algorithm for speckle denoising (MM-despeckle)

The MM based MLE algorithm involves the following steps:

- 1. Initialize x_0_ = y
- 2. *k^th^* iteration Minimize *M′* (x; x^*k*^)
- 3. x^*k*+1^ = argmin_x_ *M′*(x; x^*k*^)
- 4. If not converged: x^*k*^ = x^*k*+1^, Otherwise exit

We use gradient descent to solve step 2, so the overall problem reduces to solving an iterative gradient descent approach. We assume convergence if the change between minima of consecutive iterations x^*k*+1^ and x^*k*^ is less than 10^−10^. The method is implemented in 2D planes, but can be extended to 3D volumetric data as well.

## IV. Material and Methods

### A. Simulation

GGD speckle 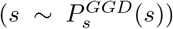 and zero mean Gaussian distributed additive noise with a standard deviation of *σ* (*n ~ N*(0, *σ*)) were simulated and used to corrupt a ground truth OCT image of a human visual cortex sample,

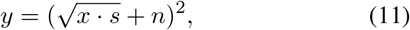

where *x* is the intensity of an arbitrary voxel in the ground truth OCT image, *y* is the measured intensity of the same voxel. We define the signal to noise ratio (SNR) as the reciprocal of *σ* in the rest of the paper. In addition we multiplied an exponential decay to simulate the light propagation in the tissue,

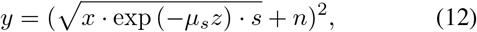

where *μ_s_* (*μm*^−1^) is the optical scattering coefficient and *z* is the double-path of light propagation in *μm*. Figure 3. shows (a) the ground truth human visual cortex tissue image that was used in our simulations, (b) a noisy image that is corrupted by GGD speckle with *α = β = γ* = 1 and additive noise with *σ* = 0.05, as described in eqn. 12, (c) the mean MM-despeckle result calculated across 100 realizations of speckle corrupted data.

**Fig. 3.**
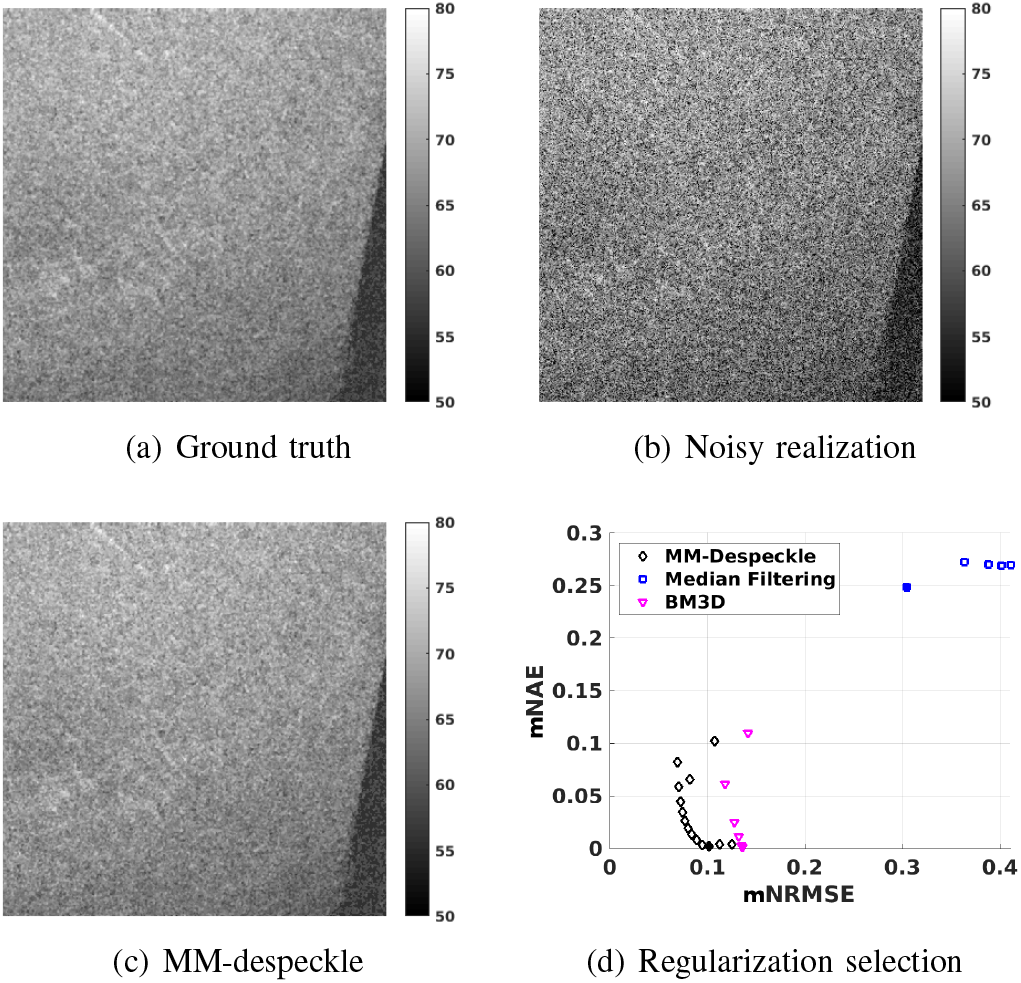
The first three images show the simulation setup - (3a.) ground truth, (3b.) one noisy image corrupted by speckle and additive noise and (3c.) The mean of MM-despeckle result calculated across 100 realizations. Figure 3d. plots mNAE versus mNRMSE plot for multiple method parameters of median filtering, BM3D and MM-despeckle methods. The point that is filled in has the lowest mNAE for each method and was used to select optimal parameters.

MM-despeckle’s performance was compared with 2D median filtering and BM3D [22] to remove speckle from 100 realizations of simulated noisy data at SNR of 5, 20 and 50. SNR here refers to the reciprocal of additive noise standard deviation *σ*. We uniformly sampled regularization parameter from 1 to 14 for MM-despeckle, window sizes from 3 × 3 to 12 × 12 for median filter and noise variance parameter from 0.0001 to 0.3 for BM3D. MATLAB 2018b medfilt2 function and BM3D MATLAB package from http://www.cs.tut.fi/foi/GCF-BM3D were used in this simulation.

We calculated normalized root mean squared error (NRMSE) with the ground truth for each despeckled image to evaluate performance,

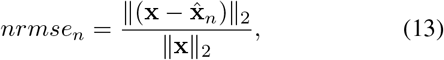

where x is the ground truth image, 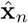 is the estimated image, *n* is an index for the noisy realization that goes from 1 to 100 and || · ||_2_ is the *ℓ_2_* norm operator.

In addition, to choose the regularization parameter we calculated mean NRMSE (mNRMSE) and mean normalized absolute error (mNAE)

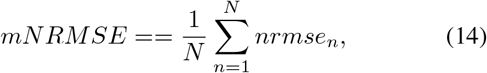

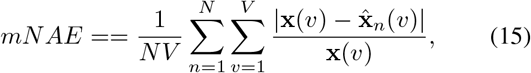

where *v* is an arbitrary voxel, *V* and *N* are the total number of voxels and noisy realizations respectively. We chose method parameter that resulted in the least mean NAE calculated across all voxels for each of the three methods. NRMSE values were compared at multiple SNRs and imaging depths.

### B. OCT imaging and speckle distribution fitting

We used a spectral domain OCT system at a center wavelength of 1300 *nm* to image a scattering phantom and various tissue structures of postmortem human brain [45]. The axial resolution was 3.5 *μm* in tissue. Spectrometer consisted of a 1024 pixel line scan camera operating at an A-line rate of 47 *kHz*. The total imaging depth was 1.5 mm, with a voxel size of 2.9 *μm*. The sensitivity of the system was 105 *dB*. We used three sets of objective lenses to test the denoising algorithm with varying lateral resolution, including a 10× air coupled objective and a 10× water immersion objective yielding a resolution of 3.5 *μm*, a 5x air coupled objective yielding a resolution of 6.5 *μm*, and a 20× water immersion objective yielding a resolution of 1.3 *μm*.

Cross-sectional OCT images were normalized by dividing the mean of intensities at each depth. The normalized intensity was fitted by generalized gamma distribution and the fitting parameters were used as inputs to the MM-despeckle optimization. In this study, we customized the fitting parameters for each experiment.

### C. Phantom Experiment

The scattering phantom was made by suspensions of monodisperse polystyrene microspheres with a refractive index of 1.57 at 1300 *nm* wavelength and a mean diameter of 1 *μm*. The solution was diluted with three concentrations, representing a scattering coefficient of 0.002 *μm*^−1^, 0.006 *μm*^−1^ and 0.01 *μm*^−1^, respectively, which roughly matched the range of scattering coefficient of gray and white matter in *ex vivo* human brain samples. The phantom samples were measured with a 10× air coupled objective, resulting in a lateral resolution of 3.5 *μm*. Each measurement consisted of a cross-section with 5000 A-lines.

The phantom image was denoised using 1D median filters with filter widths uniformly spaced from 3 (6 *μm*) to 21 (60 *μm*). The phantom image was normalized with the mean value before removing speckle with MM-despeckle. The normalization scales the range of regularization parameters. Uniform spaced regularization parameter in the range of 5000 to 50000 was found suitable for this data. Pixel-wise scattering coefficients were estimated using the approach described in [7] from the original image without speckle removal, median filtering result and the proposed method results. Error metrics mNRMSE and mNAE were calculated for the estimated scattering coefficients. Optimal regularization parameters for each of the speckle removal methods were selected as the one with the least mNAE. NRMSEs of the estimated coefficients of all three were compared.

### D. Tissue experiment and validation

Four blocks of sample including hippocampus, visual cortex, cerebellum, and brainstem were obtained from a postmortem human brain at the Massachusetts General Hospital Autopsy Suite. The samples were fixed with 10% formalin for two months. The postmortem interval did not exceed 24 hr.

The hippocampal tissue was imaged in a 10×10×2 *mm* block using a 10x water immersion objective. The block was scanned in consecutive tiles with 90% overlap. The tiles were stitched together to form a whole surface and serial sectioning was used to cover the entire depth [46]. The optical resolution was 3.5 *μm* isotropic. One imaging tile covers a volume of 1.5×1.5×1.5 mm, with an isotropic voxel size of 2.9 *μm*. The extensive overlap between adjacent tiles offers spatial averaging for denoising. With 90% overlap, a single tile can be averaged up to 100 times with an expectation of 10 fold reduction in speckle contrast. We have verified on the data that the speckle patterns are decorrelated between adjacent tiles. As a result, the stitched image serves as a reference to evaluate the performance of the MM-despeckle algorithm. In addition, the human cerebellum, visual cortex, brainstem and individual neurons in the cortex were imaged with a resolution of 6.5 *μm*, 3.5 *μm*, 3.5 *μm*, and 1.3 *μm*, respectively, using a 50% overlap between adjacent tiles. Unlike in our simulations or phantom experiment, human tissue data was processed without normalization. Therefore the scaling of regularization parameters was observed to be vastly different and in the range of 0.001 to 0.1.

## V. Results

### A. Simulation results

Figure 3d. plots the mNAE and mNRMSE of the median filtering, BM3D and MM-despeckle results at the surface (depth 0 *μm*) of the tissue with SNR=50. Each point in scatter plot in 3d. corresponds to a specific method parameter. The marker is filled in with color for the case with least NAE, and the corresponding parameter was chosen as optimal. While both MM-despeckle and BM3D reduce mean NAE close to zero, median filtering suffers from higher mean NAE of 0.25 (25 %). Optimal median filter size of 3×3, BM3D noise variance of 0.2 and MM-despeckle regularization of 5 resulted in least mNAE and were set for the rest of the error comparisons.

Figures 4. summarizes the simulation results at three different SNRs (50, 20 and 5). Boxplots show the distribution of NRMSE across 100 noisy realizations at the tissue surface (0 *μm*). Figure. S2 in the supplementary section plots the NRMSE errors for the same three SNRs but at 200 *μm* optical depth, that is typical scale in ex vivo OCT imaging. MM-despeckle demonstrates the lowest NRMSE among all the three methods at both depths and at all three SNRs. Median filtering has the highest NRMSE with a average error of around 30% at both depths. MM-despeckle and BM3D have NRMSE average in the range 9%-10% and 13%-14% respectively. MM-despeckle demonstrates an improvement of 4% in the errors at all SNRs and depths compared to BM3D. The bias in BM3D estimates can potentially be due to incorrect likelihood assumption. In contrast MM-despeckle uses a more accurate likelihood formulation, regularizes with a quadratic function that matches the additive Gaussian noise likelihood and penalizes similarity across two adjacent voxels.

**Fig. 4.**
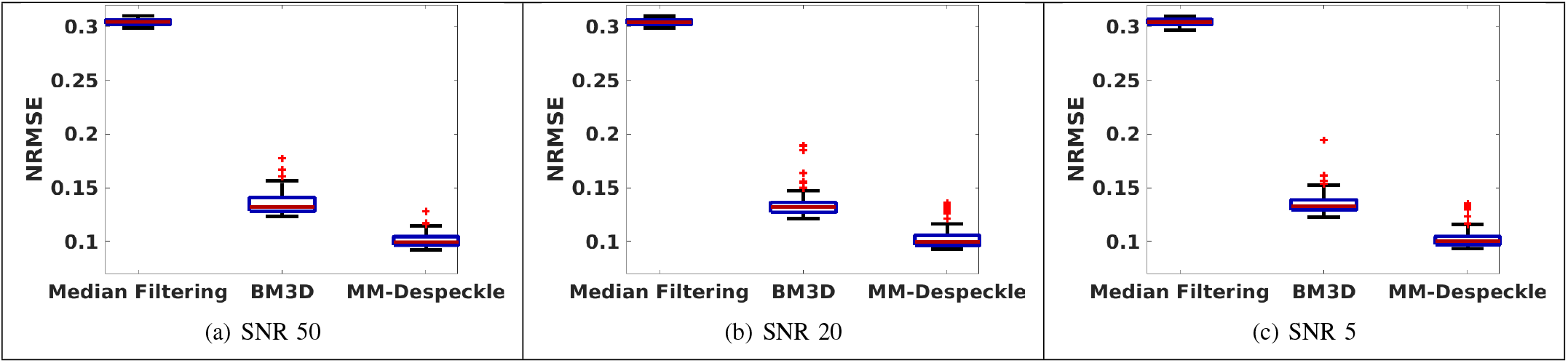
The figures plots NRMSE values across multiple noise realizations for median filtering, BM3D and MM-despeckle at SNRs 50, 20 and 5. MM-despeckle consistently demonstrates the lowest errors across SNRs.

Median filtering has the least variance across SNRs compared to the other two methods. We observe that the variance of BM3D and MM-despeckle increases with reducing SNR. This suggests that median filtering optimizes the variance strongly and suffers from large bias error, while the other two approaches reduce the error in the bias but suffer from marginally higher variance when SNR drops.

### B. Phantom experiment results

Figure 5. compares the ground truth scattering coefficient (*μ_s_*) with the scattering coefficient estimated from phantom data without removing speckle, the median filtering method (6 *μm* and 60 *μm* filter sizes), and the proposed MM-despeckle method. The approach in [7] was used to estimate the coefficients. We show the comparisons for a phantom with ground truth coefficient of 0.006 *μm*^−1^ (*α* = 1.21, *β* = 1.35, *ξ* = 0.85) in Fig. 5a. Two more phantom comparisons with coefficients 0.01 *μm*^−1^ (*α* = 1.14, *β* = 1.20, *ξ* = 0.92) and 0.002 *μm*^−1^ (*α* = 1.24, *β* = 1.42, *ξ* = 0.82) are additionally shown in Fig S2. in the supplementary section. The GGD parameters reported here were the result of fitting the distribution to the mean normalized phantom data using [44]. Median filter with the largest filter size of 21 points (60 *μm*) and MM-despeckle regularization parameter of 24000 (3 *μm*) resulted in the lowest mNAE for each of the methods. Figures 5a, 5b. and 5c. plots the scattering coefficient estimated from individual A-lines of different methods for the phantom data. For the median filtering method we plot the result of using a filter of size 3 (6 *μm*) and the optimal filter (21, i.e. 60 *μm*). Filter size of 6 *μm* is more typical of what is used with real data. The scattering coefficient that was estimated from the mean of 5000 A-lines converged to the theoretical value with high accuracy for all three phantoms and is marked as the measured ground truth in the plots. Figure 5b. plots the NRMSE of the coefficient estimates calculated for the three phantoms shown in 5a. and S2. and two additional ones with scattering coefficients 0.004 *μm*^−1^ (*α* = 1.17, *β* = 1.29, *ξ* = 0.85) and 0.008 *μm*^−1^ (*α* = 1.13, *β* = 1.19, *ξ* = 0.92). MM-despeckle demonstrates the lowest NRMSE compared to noisy and the two median filtered results.

**Fig. 5.**
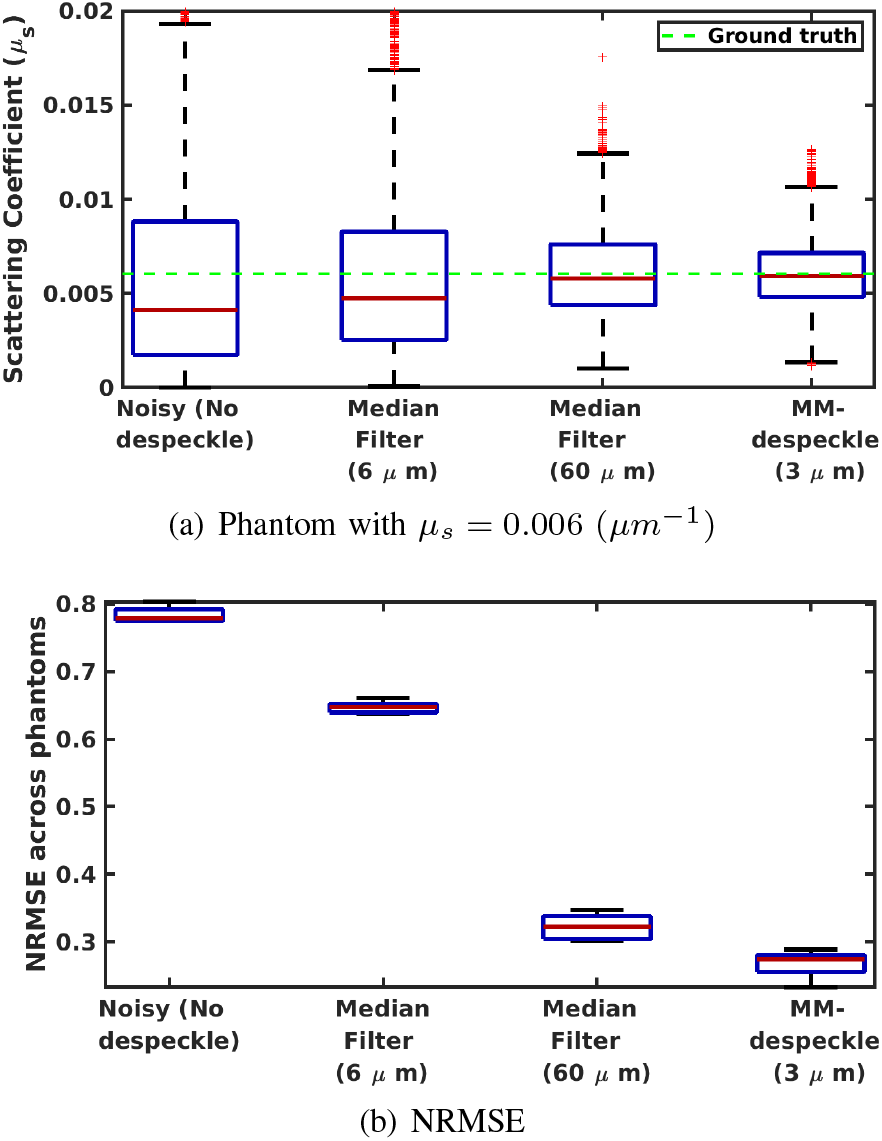
Figure 5c. compares the accuracy of the scattering coefficient estimate of MM-despeckle and median filtering with 6*μm* and 60 *μm* filter sizes in a uniform phantom with optical property 0.006 *μm*^−1^. 5d. plots the NRMSE calculated across phantoms

Compared to the original data and the 6 *μm* median filter, MM-despeckle consistently demonstrates lower errors and fewer outliers in the coefficient estimation results of all three phantoms. MM-despeckle either matches or is better in accuracy than the 60 *μm* median filter that uses all 21 points. However, the 60 *μm* filter size is unsuitable to use in biological microstructure imaging because it will smooth boundaries and features that are smaller than 60 *μm*, which is commonplace. On the contrary, MM-despeckle smooths within a 3 *μm* radius and is therefore more applicable for preserving microstructures.

### C. Hippocampus imaging experiment

In this section we present MM-despeckle results of the human hippocampus imaged by OCT. We calculated the speckle contrast (std deviation/mean intensity) for a region of the hippocampus with several regularization parameters uniformly spaced from 0 to 0.01. Although speckle contrast keeps decreasing with regularization, increasing the regularization also smooths the image thereby blurring the edges. Keeping this trade-off in mind, we selected the regularization parameter based on an optimal speckle contrast set by the experimental result of overlapping tiles.

Figure 6a plots the rate of decrease in speckle contrast with increasing number of averages from overlapping tiles. With 7 averages, the reduction rate drops to 5%, beyond which we consider that the speckle contrast does not decrease much anymore. We choose the speckle contrast corresponding to 7 averages (= 0.25) to be optimal. Next we found that the corresponding regularization parameter of 0.007 leads to the optimal speckle contrast in the output. The variation of speckle contrast with regularization parameter is shown in Figure 6b. We used the same regularization parameter hereinafter for all the tissue imaging results. We also set all the three GGD parameters to 1 for all human tissue experiments based on the average fit from our phantom data.

**Fig. 6.**
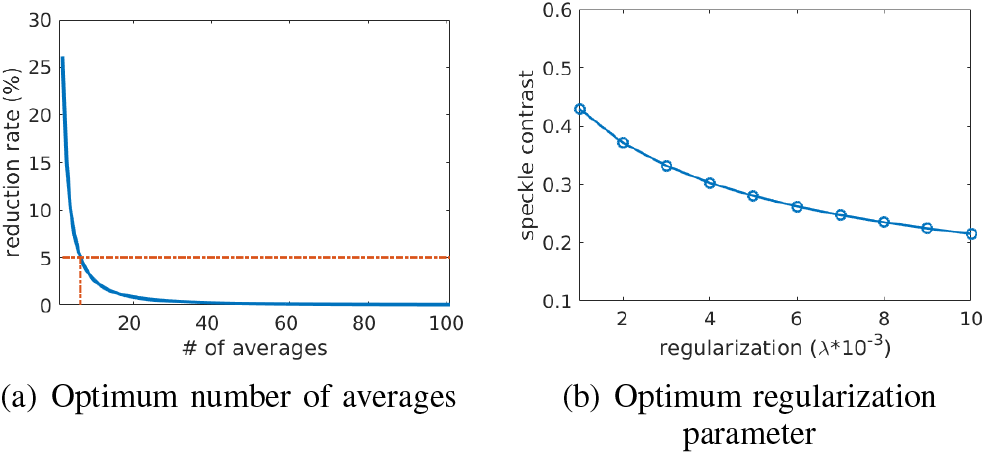
Speckle contrast based selection of (a) number of averages and (b) regularization parameter.

We compared the MM-despeckle results with BM3D and median filtering. For medan filterin g, filter size of 3 × 3 was chosen because at 5×5 we start observing blurring in our results. Similarly, for BM3D, noise variance of 0.2 was chosen as at 0.3 we start observing blurred microstructure. Figure 7. shows a 1.5 *mm* × 1.5 *mm* slice of the hippocampus tissue. Figure 7a. is the original hippocampus image without denoising, 7b. is the MM-despeckle result with the regularization parameter set to 0.007, 7c. and 7d. are results of median filtering with filter sizes 3× 3 and 5× 5 respectively, 7e. and 7f. are the BM3D denoising results with noise variance of 0.2 and 0.3 respectively.

**Fig. 7.**
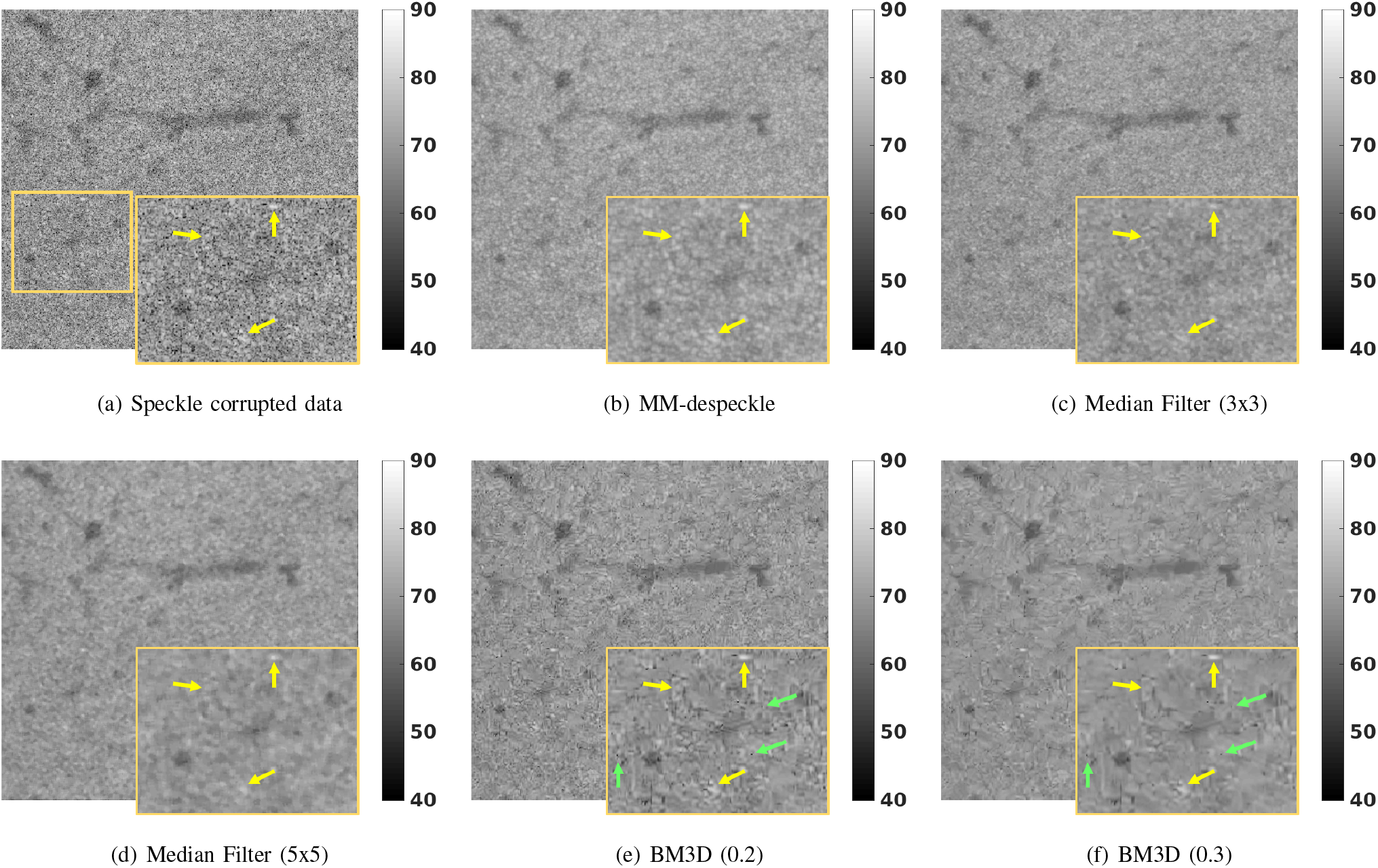
The figure compares MM-despeckle, BM3D and Median filtering methods qualitatively applied to remove speckle from human hippocampus OCT data. MM-despeckle preserves small features better than median filtering and reduces speckle better than BM3D due to accurate modeling.

MM-despeckle removes the speckle and recovers the tissue contrast successfully. Both BM3D results have holes in their images (marked with green arrows) due to uncorrected speckle as it does not model speckle accurately. In addition, BM3D suffers from blurring the anatomy unnaturally because at high resolution it is unable to match similar regions well. The 5 × 5 median filter result blurs the anatomy as expected. The 3 × 3 median filter removes speckle well but suffers in the regions of small features with high intensities marked by yellow arrows. This can be problematic for imaging small tissue structures such as vessels or amyloid deposits on the tissue that show up in images as high intensities.

MM-despeckle is demonstrably a better option than both BM3D and median filtering because it corrects speckle well, blurs the anatomy less and retains small structures successfully (see yellow arrows). In addition, for this dataset BM3D performed worse than median filtering as it was unable to find similar blocks. On the contrary, in our simulations where the data contained more similar blocks, BM3D outperformed median filtering. Therefore the performance of BM3D as compared to median filtering is data dependent. However, MM-despeckle consistently outperformed both BM3D and median filtering for both datasets.

### D. MM-despeckle minimizing acquisition time

It is generally challenging to assess the performance in tissue data where we do not know the ground truth of OCT intensity or scattering coefficient. For this reason we acquired data from a human hippocampus with 90% overlapping tiles, which means that every voxel was acquired 100 times with independent speckle patterns. The speckle reduction rate with averaging was presented in figure 6a. Overall speckle reduction ratio was calculate by taking the ratio of the speckle contrast with 100 averages to no average. Taking averages of the 100 measurements reduced the speckle by a factor of 10 and provided us with a good reference image.

Figure 8. qualitatively compares (a) the original speckle corrupted image stitched without averaging, (b) MM-despeckle applied to the noisy image in 8a. and (c) the 100-averaged reference image of the hippocampus sample. MM-despeckle image successfully removes the speckle in the original data. NRMSE was calculated for a region at the center of the sample of size 6 mm × 6 mm × 0.25 mm for no-average and MMde-speckle result with the reference image. NRMSE reduced by 50 percent for MM-despeckle (NRMSE=0.76) compared to the no-average image (NRMSE=1.37). The reduction in error by 50 percent for a single image means that we will not require as many averages to obtain a result that is comparable to the reference image. This result demonstrates the promise of MM-despeckle to reduce the overall acquisition time by reducing the amount of spatial averaging necessary for the imaging experiment.

**Fig. 8.**
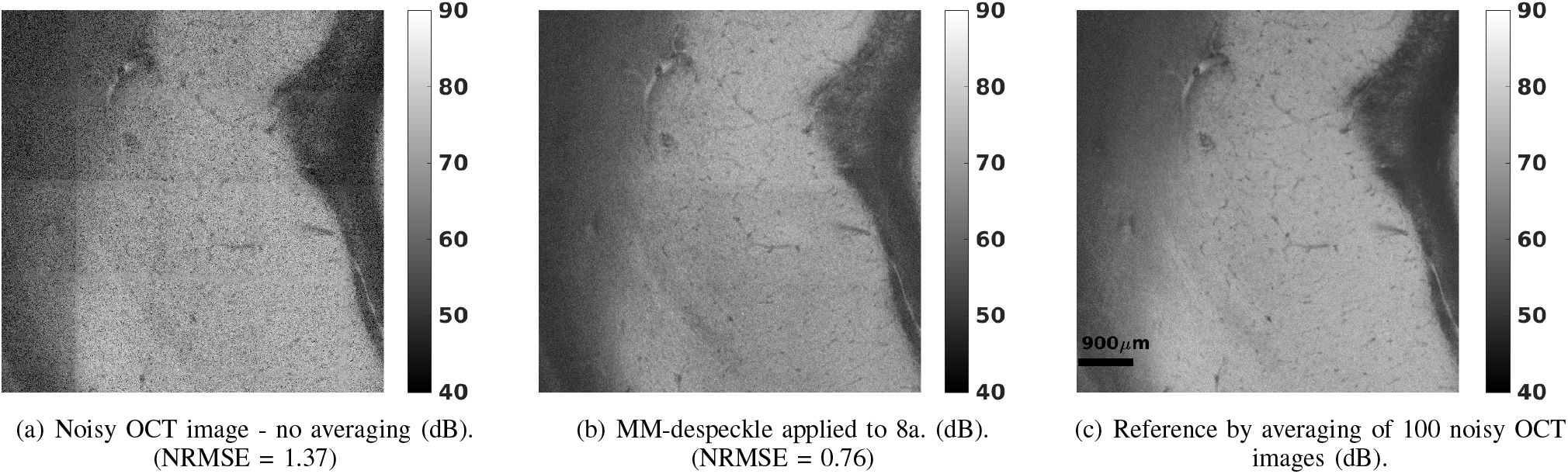
The figure compares noisy human hippocampus tissue OCT data, the result of MM-despeckle applied to the noisy OCT data and the 100-averaged reference image. MM-despeckle removes speckle, reduces the NRMSE and requires reduced spatial averaging.

### E. Generalizability of MM-despeckle results across brain structures and imaging resolutions

Figure 9 demonstrates the generalizability of MM-despeckle to remove speckle with various structures and across multiple OCT resolutions. Figures 9a, 9c, and 9e are the original OCT images of the human cerebellum, visual cortex, and brainstem respectively. The cerebellum image covers 2.9 *mm* × 2.9 *mm* at 6.5 *μm* voxel resolution while the visual cortex and the brainstem images cover 1.8 *mm* × 1.8 *mm* at 3.5 *μm* voxel resolution. Figures 9b, 9d and 9f are result of removing speckle using MM-despeckle for the three different tissue data. MM-despeckle successfully removes speckle across multiple tissue types and across two resolution scales without blurring the anatomy. Specifically we observe small features such as the high intensity deposits in the cerebellum more clearly visible after the correction. The same regularization parameter of 0.007 was used for all three cases further demonstrating the robustness and generalizability of the MM-despeckle method across the different tissue types and OCT imaging resolutions.

**Fig. 9.**
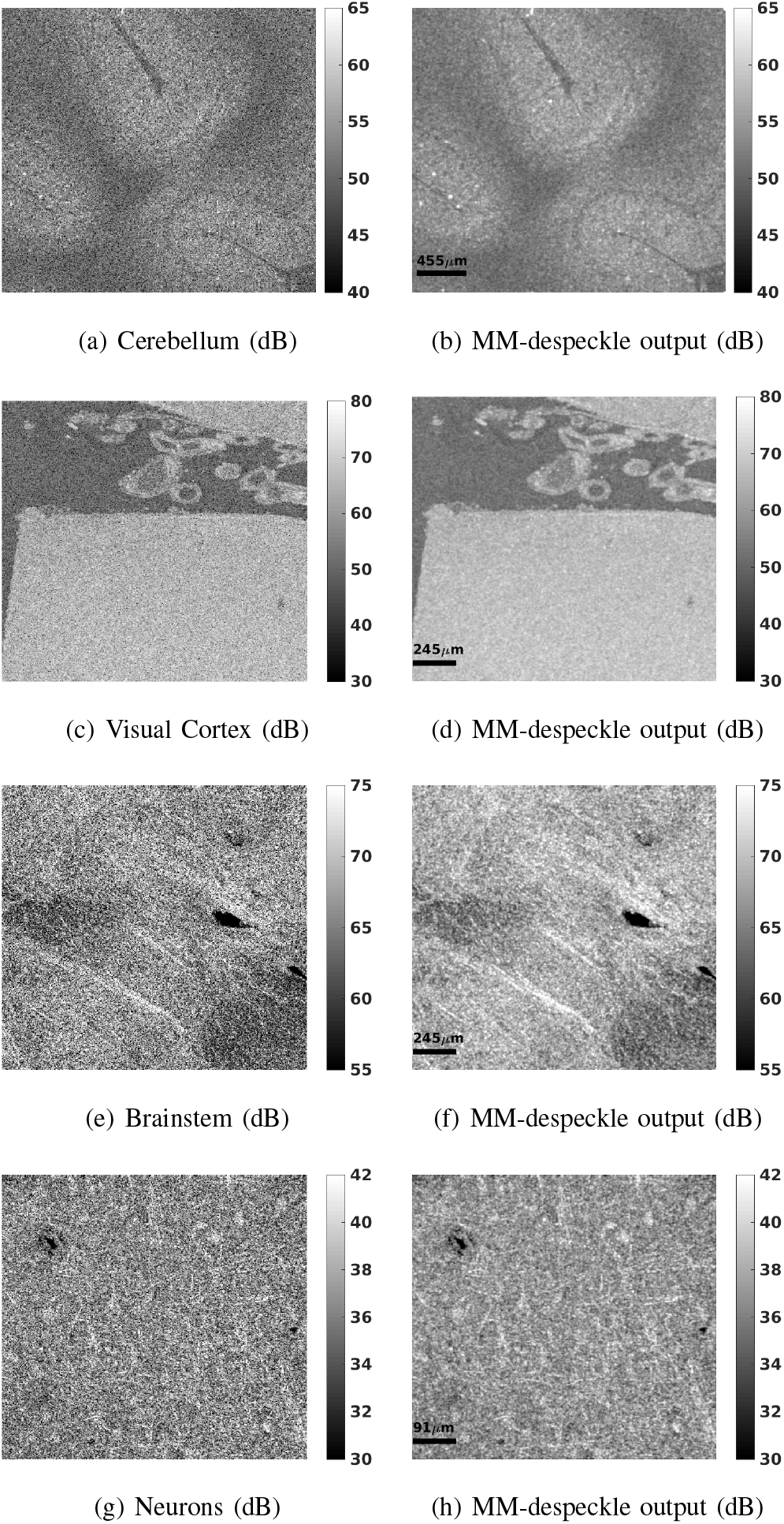
The figures demonstrate the generalizability of MM-despeckle across various human microstructure tissue and varying resolutions. Figures 9a-c correspond to tissue from visual cortex, cerebellum and brainstem acquired at 3 *μm* isotropic resolution. Figure 9d. corresponds to an image of neurons from the visual cortex imaged at 1.5 *μm* isotropic resolution.

Figure 9g and 9h are the original and MM-despeckle method result for optical coherence microscopy (OCM) images of neurons in the cortex with 0.67 *mm* × 0.67 *mm* image size at 1.3 *μm* voxel resolution. The regularization parameter was set the same as above tissue types. MM-despeckle removes speckle successfully without blurring thereby improving the contrast of the neurons with the background. The result further demonstrates that the proposed MM-despeckle method is applicable to OCM images at single cell resolution without the need to adjust regularization parameters and therefore is strongly generalizable across multiple imaging resolutions.

## VI. Discussion

In this work we proposed a new majorize-minimize-based optimization method called MM-despeckle to remove generalized gamma distributed multiplicative speckle noise from OCT images. There are three major contributions in this work.

1. We observed a generalized gamma distribution for characterizing the statistics of speckle based on real imaging data and built a statistical model to remove the speckle and restore the microstructure images of human brain samples. Particularly, the parameters of the distribution function were tuned for each experiment and the model can be simplified to gamma or negative exponential distribution depending on OCT system and tissue properties.
2. An optimization framework was proposed to solve the non-convexity of the generalized gamma P-NLL problem. Although applied to OCT in the current study, this theoretical framework is applicable to other imaging modalities contaminated by speckle noise.
3. The MM-despeckle significantly reduces the acquisition time otherwise 10-90 times longer in ex vivo OCT imaging, due to the requirement of extensive averaging to achieve satisfactory CNR.

MM-despeckle minimizes a P-NLL based cost function that is standard for statistical estimation problems. We have used a quadratic smoothness-based spatial regularization for our results. However, the framework itself can be seamlessly integrated with other convex regularization functions such as total variation [19] and/or wavelet transformation based functions [47] that have been used in other speckle removal applications and that have analytical or numeric way to calculate gradients. We also demonstrated the effectiveness of using speckle contrast changes to select the regularization parameter for tissue data where we do not have the ground truth. While this worked well for our microstructure application, for applications where speckle contrast is not a suitable criteria other regularization parameter selection methods such as those in [48] can also be utilized.

The theoretical novelty of our approach is the proof of non-convexity of the generalized gamma NLL. As is the case with all non-convex optimization problems, the proposed MM-despeckle method might get trapped in a local rather than global minimum. For all the results shown, we initialized MM-despeckle with the original image because doing so in our simulation and phantom experiments resulted in least error compared to other methods. Moreover, for the simulation initializing MM-despeckle with the output of median filtering resulted in a local minimum that suffered from bias similar to that of median filtering. This observation suggested that the median filtering output is close to a plausible suboptimal local minimum solution and hence not suitable for initializing our proposed method. While our choice of noisy image-based initialization has proved to be robust for all our experiments including real tissue data, this by no means guarantees a theoretical global minimum. Global optimum search strategies such as using multiple initializations can be incorporated into our approach to further improve the results.

A common next step after removing speckle for OCT images is to calculate the scattering coefficients. We demonstrated the improvement in the accuracy of estimating the coefficients with MM-despeckle in our phantom experiment. We have extended MM-despeckle to jointly remove speckle and estimate the coefficients in one step and presented the initial results in [40]. This extension avoids a two step process of first removing the speckle and then estimating the coefficient, and instead combines the two into a single procedure. We demonstrated promising results that improved accuracy of the coefficient estimation even further and will be performing detailed analysis in future work.

Lastly, while the examples in this paper primarily focus on OCT imaging, the approach is relevant to several other applications such as RADAR, SONAR and other optical imaging modalities where generalized gamma distribution-based speckle noise has been shown to be problematic. MM-despeckle can be applied to those settings as well.

## VII. Conclusion

We proposed a new method called MM-despeckle to remove speckle from OCT images. The approach optimizes regularized generalized gamma distributed NLL cost function iteratively. We carried out simulation, phantom and tissue experiments to demonstrate the usefulness, generalizability and improved performance of the method compared to the state of the art. Future work focuses on jointly estimating scattering coefficient along with removing speckle.

## Notes

\# In addition, B. Fischl has a financial interest in CorticoMetrics, a company whose medical pursuits focus on brain imaging and measurement technologies. B. Fischl’s interests were reviewed and are managed by Massachusetts General Hospital and Partners HealthCare in accordance with their conflict of interest policies.

Support for this research was provided in part by the BRAIN Initiative Cell Census Network grant U01MH117023, the National Institute for Biomedical Imaging and Bioengineering (P41EB015896, 1R01EB023281, R01EB006758, R21EB018907, R01EB019956, R00EB023993, U01EB026996), the National Institute on Aging (1R56AG064027, 1R01AG064027, 5R01AG008122, R01AG016495), the National Institute of Mental Health, the National Institute of Diabetes and Digestive and Kidney Diseases (1-R21-DK-108277-01), the National Institute for Neurological Disorders and Stroke (R01NS0525851, R21NS072652, R01NS070963, R01NS083534, 5U01NS086625, 5U24NS10059103, R01NS105820), and was made possible by the resources provided by Shared Instrumentation Grants 1S10RR023401, 1S10RR019307, and 1S10RR023043. Additional support was provided by the NIH Blueprint for Neuroscience Research (5U01-MH093765), part of the multi-institutional Human Connectome Project.

### Competing Interest Statement

The authors have declared no competing interest.

