## Supplementary Material for "A novel algorithm to optimize generalized gamma distributed multiplicative noise with implications on speckle removal from OCT images"

This supplement contains additional material to complement the results presented in the main body of the paper. Majorant derivations for gamma and negative exponential distributions are derived in Sec. I., while supplementary figures are found in Sec. II.

### I. Majorants for gamma and negative Exponential distributions

Generalized gamma distribution (GGD) (Eq. 3) simplifies to the gamma distribution (GD) when $\xi = 1$ and to the negative exponential distribution (NED) with $\alpha = \beta = \xi = 1$,

$$\begin{aligned} P_S^{GD}(s;\alpha,\beta) &= P_S^{GGD}(s;\alpha,\beta,1) \\ &= \frac{\beta^\alpha}{\Gamma(\alpha)} s^{\alpha-1}\exp(-\beta s), \end{aligned} \tag{1}$$

$$\begin{aligned} P_S^{NED}(s) &= P_S^{GGD}(s;1,1,1) \\ &= e^{-s}, \end{aligned} \tag{2}$$

where $\alpha$ and $\xi$ are the shape parameters and $\beta$ is the rate parameter of GGD.

#### A. Gamma distribution

We derive analytical expressions of joint likelihood distribution, P-NLL cost and majorant of GD by substituting $\xi = 1$ in equations 7, 8 and 9 in the main text respectively. The three quantities reduce to:

GD joint likelihood:

$$P_{Y|X}(\mathbf{y}|\mathbf{x}) = \beta^{M\alpha}\prod_{m=1}^{M}\left[y_m^{\alpha-1}x_m^{-\alpha}\exp^{-y_m/x_m}\right]. \tag{3}$$

GD P-NLL cost:

$$\begin{aligned} \hat{\mathbf{x}} &= \arg\min_{\mathbf{x}} -\log(P_{Y|X}(\mathbf{y}|\mathbf{x}) + \lambda R(\mathbf{x}), \\ &= \arg\min_{\mathbf{x}} \sum_{m=1}^{M}(\alpha\log(x_m) + \beta\frac{y_m}{x_m}) + \lambda R(\mathbf{x}). \end{aligned} \tag{4}$$

GD majorant:

$$M(\mathbf{x};\mathbf{x}^i) = \sum_{m=1}^{M}\alpha\left(\frac{x_m}{x_m^i} + \log(x_m^i) - 1\right) + \beta\frac{y_m}{x_m}. \tag{5}$$

#### B. Negative exponential distribution

We further reduce the equations in I-A by substituting $\alpha = \beta = 1$ in each of the equations and derive the expressions for the NED case:

NED joint likelihood:

$$P_{Y|X}(y|x) = \prod_{m=1}^{M}\left[\exp^{-y_m/x_m}/x_m\right]. \tag{6}$$

NED P-NLL cost:

$$\begin{aligned} \hat{\mathbf{x}} &= \arg\min_{\mathbf{x}} -\log(P_{Y|X}(\mathbf{y}|\mathbf{x}) + \lambda R(\mathbf{x}), \\ &= \arg\min_{\mathbf{x}} \sum_{m=1}^{M}(\log(x_m) + \frac{y_m}{x_m}) + \lambda R(\mathbf{x}). \end{aligned} \tag{7}$$

NED majorant:

$$M(\mathbf{x};\mathbf{x}^i) = \sum_{m=1}^{M}\frac{x_m}{x_m^i} + \log(x_m^i) - 1 + \frac{y_m}{x_m}. \tag{8}$$

### II. Supplementary Figures

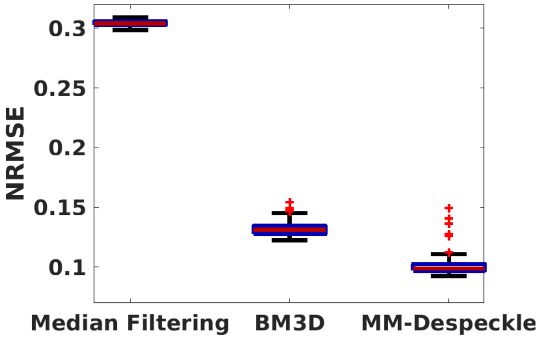


(a) SNR 50

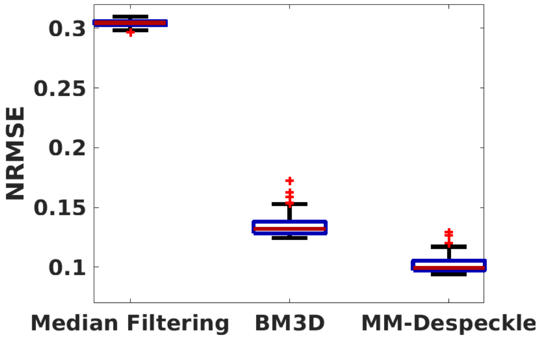


(b) SNR 20

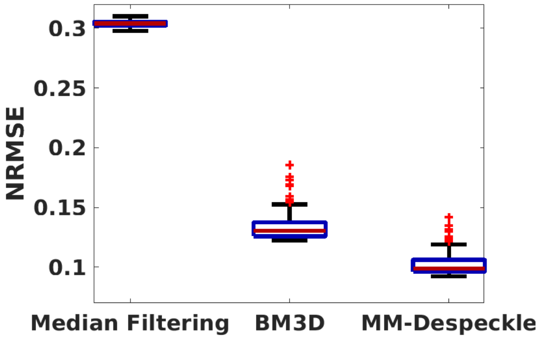


(c) SNR 5

Fig. S1: The figures plots NRMSE values across multiple noise realizations at 200 $\mu m$ depth for median filtering, BM3D and MM-despeckle at SNRs 50, 20 and 5. MM-despeckle consistently demonstrates the lowest errors across SNRs and depths.

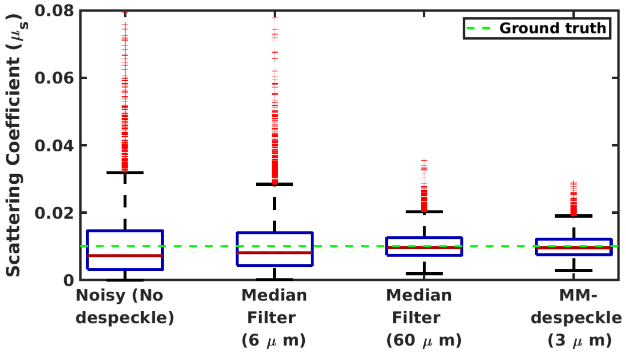


(a) Phantom with $\mu_s = 0.01$ $(\mu m^{-1})$

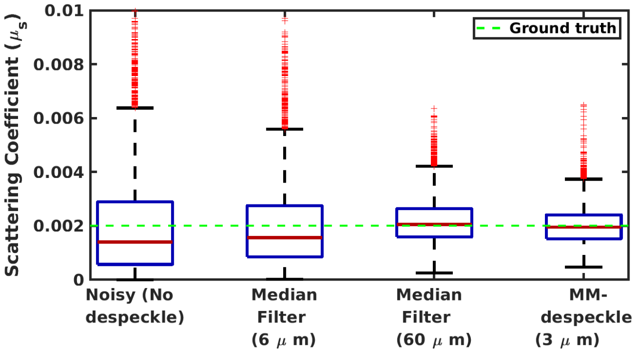


(b) Phantom with $\mu_s = 0.002$ $(\mu m^{-1})$

Fig. S2: The figure compares the accuracy of the scattering coefficient estimate of MM-despeckle and median filtering with 6$\mu m$ and 60 $\mu m$ filter sizes in two uniform phantoms with optical property - (5a.) 0.01 $\mu m^{-1}$ and (5b.) 0.002 $\mu m^{-1}$.
